# Measurement-Induced Decision Loss: A feasible-set theory of microbiome intervention under observation loss

**DOI:** 10.64898/2026.10.06.757205

**Authors:** Javad Aminian-Dehkordi, Mohammad Mofrad

## Abstract

Microbiome measurements are commonly judged by how faithfully they reconstruct an underlying biological profile, using taxonomic similarity, prediction accuracy, or compositional distance. Yet faithfulness and usefulness are not the same: a distorted measurement may still support the correct intervention, while a small error can be decisive when it falls near the boundary between two treatment choices. We therefore evaluate measurement quality by whether information loss changes the decisions and outcomes that remain attainable. We treat the intervention-relevant state of the gut community as a hidden quantity seen only through an imperfect, information-losing measurement, mapped by a decision rule to a graded intervention. Each decision rule is evaluated by net host benefit and ecological margin, generating a feasible set of attainable outcome pairs. We define the measurement-induced decision loss (MIDL) as the maximum net host benefit lost under imperfect measurement relative to perfect observation. For scalar affine information loss, increasing degradation is a Blackwell garbling; under post-processing closure, more informative measurement weakly expands the feasible set and cannot reduce optimal host benefit. In the binary-state model, MIDL increases linearly while an observation-dependent decision rule remains optimal and plateaus once the optimal policy becomes signal-independent. We extend the framework to finite states, state-specific loss, multi-segment stool aggregation, and ecological-margin constraints. The framework distinguishes measurement faithfulness from decision usefulness and provides a principled basis for comparing stool and richer measurements by the interventions and trade-offs they support.

## 1 Introduction

Microbiome-guided intervention begins with a measurement, and in most human gut studies, that measurement is obtained from stool because stool collection is non-invasive, repeatable, and scalable. However, it is an endpoint sample, not a direct observation of the gut microbiome. The gastrointestinal tract contains spatially distinct physicochemical niches, and microbial composition and metabolic activity vary across regions and over time [Donaldson et al., 2016, Folz et al., 2023]. Direct sampling of intestinal contents has further shown that microbial and omic profiles within the gut can differ substantially from what appears in stool profiles [Shalon et al., 2023]. Stool should therefore be viewed as a transformed, spatially averaged signal whose adequacy depends on the scientific or clinical task for which it is used.

Most assessments of that adequacy ask how faithfully a stool profile reproduces an intestinal quantity: how well it correlates, classifies a state, or predicts an outcome. These are reasonable questions, but predictive accuracy alone does not determine the usefulness of a measurement for intervention. A distortion that is small on average may be consequential if it changes the intervention tier selected near an intervention boundary, whereas a visibly imperfect measurement may be sufficient when all plausible corrections lead to the same decision. Measurement quality is therefore partly a property of the downstream decision problem, including the available intervention tiers, their benefits and burdens, and the distribution of states in the target population.

Decision theory provides a natural language for making this distinction. In Blackwell’s comparison of experiments, one measurement is more informative than another when the latter can be generated by post-processing, or garbling, the former. For a decision-rule class closed under such post-processing, the more informative measurement cannot yield a lower optimal expected utility [Blackwell, 1953]. This ordering is stronger than a comparison based on a single prediction metric because it evaluates information through the decisions that the signal can support.

A related feasible-set perspective appears in information-design analyses that characterize attainable pairs of stakeholder utilities under alternative information structures [Bergemann et al., 2015]. Probabilistic masking can likewise alter the geometry of an attainable utility region [Fallah et al., 2024]. The connection used here is structural rather than literal: latent states are intervention-relevant community states, signals are microbiome measurements, and outcomes are net host benefit and ecological margin.

For microbiome intervention, however, a single host-benefit objective may be insufficient. An intervention may improve the expected host benefit while imposing treatment burden, substantially perturbing the microbial community, or reducing the ecological reserve (safety buffer) remaining after intervention. We refer to this modeled reserve as the ecological margin. Ecological resistance– the degree to which composition or function withstands disturbance–and resilience–the extent or rate of recovery after disturbance–are distinct properties [Allison and Martiny, 2008, Shade et al., 2012, Relman, 2012]. Neither is necessarily captured by host benefit alone. A useful theory should therefore characterize not only the maximum attainable net host benefit, but also the set of host-benefit–ecological-margin outcomes supported by a given measurement channel.

Here we develop a finite-state, decision-theoretic framework for this purpose. The intervention-relevant condition of the gut microbiome is represented by a latent categorical state with a population prior. In the ordered biological specialization, higher-ranked states support the benefits of increasingly intensive intervention tiers; the ordering need not represent increasing health or increasing dysfunction. Stool-derived measurements provide noisy measurements about this state through a row-stochastic measurement channel. A decision rule (or policy) maps the observed signal to an intervention. Each state–intervention pair has a net host-benefit utility and an ecological-margin utility. The central object is the feasible set of attainable expected utility pairs, which separates the information supplied by the measurement from the trade-offs induced by the selected rule.

Within this framework, we define measurement-induced decision loss (MIDL) as the loss in maximum attainable expected net host benefit relative to perfect observation, given a specified prior belief, utility model, intervention menu, and admissible policy class. MIDL measures the opportunity lost through imperfect measurement: larger values indicate greater loss, and zero indicates no loss for the specified decision problem. We use a scalar affine channel to obtain an explicit Blackwell garbling order and apply standard information-comparison results to the feasible outcome set. We then establish closure of the convexified monotone policy class for these scalar kernels, solve the binary policy-switch thresholds and MIDL in closed form, and extend the construction to finite states and state-specific information loss.

We further develop two extensions that connect the abstract channel to biologically motivated measurement and safety questions. First, a multi-segment aggregation identity shows that replacing heterogeneous intestinal regions with a single average measurement blur introduces a correction term equal to a weighted blur–composition covariance. Second, we impose ecological admissibility by requiring interventions to satisfy a minimum ecological margin, thereby restricting the feasible utility region to decision rules that satisfy the modeled ecological criterion.

The aim is to provide a mathematical foundation for evaluating when stool measurements are sufficient to support intervention decisions, when additional intestinal measurements can improve intervention value, and why microbiome measurement technologies should be evaluated according to the intervention outcomes they make feasible.

## 2 Model primitives

### 2.1 Latent states, interventions, and utilities

Let *z ∈* Δ^*G−*1^ denote the functional-guild composition of an individual microbial community, with *G* biological guilds. A prespecified mapping *q* : Δ^*G−*1^*→ {*0, …, *K −* 1*}* defines the intervention-relevant state *X* = *q*(*z*). Let *ϕ*_*k*_ = *P* (*X* = *k*) and *ϕ ∈* Δ^*K−*1^ denote state prevalence in the target population. Thus, *z* describes a within-community composition, whereas *ϕ* is a population prior over categorical states.

For the analytical constructions, we use *K* candidate interventions *j ∈ {*0, …, *K −* 1*}* and *K* signal categories. Equal cardinalities are a modeling convenience, not a requirement of the feasible-set principle. Let *U*_H_(*j, k*) and *U*_E_(*j, k*) be finite state–intervention utility tables. Host utility is expressed as incremental net benefit relative to a specified conservative baseline. The state mapping is chosen to retain the features required for these modeled payoffs; any decision-relevant variation omitted by the coarse-graining requires a richer state model. No ordering of states, interventions, signals, or utilities is required for the general feasible-set construction.

### 2.2 Measurement operators and decision rules

A measurement operator is a row-stochastic matrix T *∈*ℝ^*K×K*^, where *P* (*O* = *o* | *X* = *k*) = T_*ko*_. The joint distribution of the latent state and observed signal is

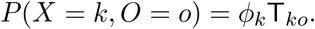

A randomized decision rule is a stochastic kernel *π*(*j* | *o*), the probability of choosing intervention *j* after signal *o*. A deterministic rule *σ* is the special case *π*(*j* | *o*) = **1** *{j* = *σ*(*o*) *}* . Restrictions on admissible rules are specified separately; monotonicity is introduced for the ordered biological specialization in Section 2.3.

For a rule *π*, the expected ecological margin and net host-benefit utilities under operator T are

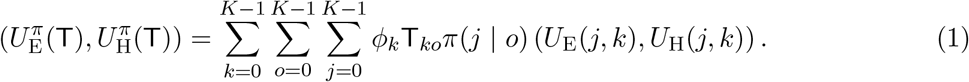

#### Definition 2.1

(Feasible outcome set). Let *P* be a class of admissible decision rules. The feasible set under measurement operator T is

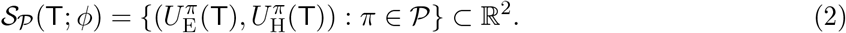

For the ordered specialization in Section 2.3, *ℳ* denotes the convex hull of deterministic monotone rules, and we write *S*_*ℳ*_(T; *ϕ*).

#### Remark 2.1

(Why the feasible set, not a single optimum?). A single optimum requires specifying a trade-off weight between host benefit and ecological margin. The feasible set separates what is attainable from how the decision-maker ultimately values the objectives. This is useful in microbiome settings where safety, stability, cost, and patient preference may be weighted differently across applications.

#### Definition 2.2

(Measurement-induced decision loss). For an admissible decision-rule class *P*, define

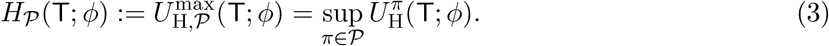

Measurement-induced decision loss is

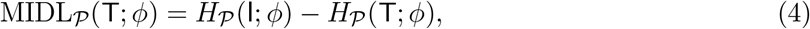

where I is perfect observation. This is the expected net host benefit forgone because the available measurement is imperfect, relative to the same decision problem under perfect observation. It can equivalently be interpreted as the value of upgrading the current channel to perfect observation, analogous to the expected value of perfect information [Howard, 1966]. It is not the benefit supplied by the existing channel relative to no information. The loss interpretation and nonnegativity require the policy-class conditions established below.

### 2.3 Ordered intervention environment as a biological specialization

For the binary and finite-state analyses, states and observed signals are ranked by their relevance to increasingly intensive intervention tiers. Tier *j* = 0 is the conservative baseline. A higher state rank means that the benefit condition for a higher tier can be met; it does not, without additional calibration, define a universal health or dysfunction scale. We use the following particular utility structure.

#### Assumption 2.1

(Ordered intervention environment). Let 0 = *b*_0_ *≤ b*_1_ *≤· · · ≤ b*_*K−*1_ and 0 = *c*_0_ *≤ c*_1_ *≤· · · ≤ c*_*K−*1_ denote gross intervention benefits and burdens. Assume that *b*_*j*_ *−c*_*j*_ is nondecreasing in *j*. Choosing tier *j* in latent state *k* yields net host utility

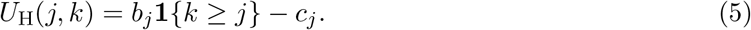

The burden *c*_*j*_ makes the cost of intervention explicit and can favor a conservative or intermediate tier when its expected benefit does not justify its burden. State-dependent effectiveness can also make personalization valuable when burdens are zero; burden is not a necessary ingredient in every finite-state decision problem. Biologically, it can represent tolerability, dietary burden, intervention risk, implementation cost, or the opportunity cost of unnecessary perturbation.

Let *ℓ*_*k*_ *≥* 0 denote the ecological reserve associated with latent state *k*, and let 0 = *δ*_0_*≤ δ*_1_*≤· · · ≤δ*_*K−*1_ denote the ecological burden imposed by the ordered intervention tiers. The ecological-margin utility is

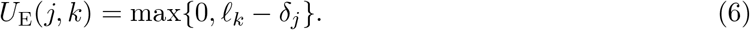

This quantity represents the ecological reserve remaining after the burden of the selected intervention. No monotonic ordering of *ℓ*_*k*_ is required unless the latent-state ordering is also intended to represent increasing ecological tolerance. Alternative calibrated ecological utilities can replace Eq. (6) without changing the feasible-set construction.

A deterministic policy is nondecreasing monotone if *o*_1_ *≤ o*_2_ implies *σ*(*o*_1_) *≤ σ*(*o*_2_). This is an admissibility convention for the ordered model, not a general property of posterior-optimal decisions. We write *ℳ* for the convex hull of nondecreasing deterministic policies. The general construction above applies to other policy classes as well.

## 3 Scalar affine observation loss and Blackwell ordering

Blackwell’s comparison of experiments supplies the standard decision-theoretic principles used in this section: stochastic garbling, feasible-set inclusion under post-processing closure, and monotonicity of optimal expected utility [Blackwell, 1953]. The explicit factorization for the selected affine family and the closure verification for its convexified monotone policy class specialize those principles to the present model. They provide the foundation for the policy thresholds, attainable trade-offs, and MIDL calculations derived in the subsequent sections.

### 3.1 Scalar affine observation channel

Let *r ∈* Δ^*K−*1^ be a reference distribution over observed signal categories. It may be uniform in a theoretical illustration or may represent a population-level signal distribution in an applied model. Let R denote the *K× K* matrix whose every row equals *r*. For *β ∈* [0, 1], define the scalar affine measurement channel

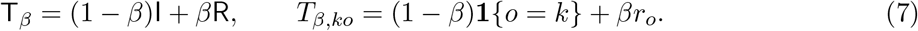

Thus, conditional on latent state *k*, the observed signal retains the state-matched category with weight 1 *− β* and is replaced by a draw from the reference distribution with weight *β*.

#### Lemma 3.1

(Row stochasticity and marginal-signal shrinkage). For every *β* ∈ [0, 1], T_*β*_ is row-stochastic. If *ϕ* is treated as a row vector, the marginal distribution of the observed signal is

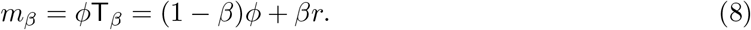

*Proof*. Each row of T_*β*_ is a convex combination of the unit vector *e*_*k*_ and the probability vector *r*, so it is nonnegative and sums to one. Moreover, *ϕ*T_*β*_ = (1 *− β*)*ϕ*I + *βϕ*R = (1 *− β*)*ϕ* + *βr*, because *r* Σ_*k*_ *ϕ*_*k*_ = 1 and every row of R equals .

Equation (8) describes shrinkage of the marginal signal distribution toward *r*. It should not be interpreted as a noiseless reconstruction of the latent state or as a within-sample microbial composition.

### 3.2 Blackwell ordering and feasible-set contraction

We compare measurement channels using Blackwell’s ordering of statistical experiments, under which a less informative channel can be generated from a more informative one by stochastic post-processing [Blackwell, 1953].

#### Assumption 3.1

(Post-processing closure). The admissible policy class is closed under signal post-processing kernels considered below. Specifically, if *π∈ P* and K is a row-stochastic kernel that maps an initial signal *o*_1_ to a post-processed signal *o*_2_, then the composed policy

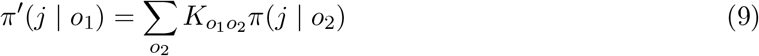

also belongs to *P*.

This condition is automatic for the class of all randomized policies. For a restricted policy class, closure must be established for the particular post-processing kernels under consideration.

#### Proposition 3.1

(Scalar affine channels are Blackwell ordered). Let 0 *≤β*_1_ *≤β*_2_ *≤*1. If *β*_1_ *<* 1, then 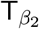 is a garbling of 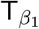. Specifically,

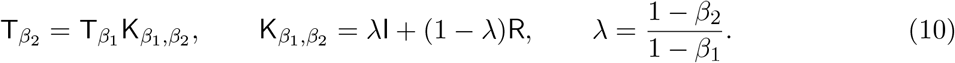

If *β*_1_ = 1, then necessarily *β*_2_ = 1, and the two channels coincide.

*Proof*. For *β*_1_ *<* 1, the ordering *β*_1_ *≤ β*_2_ implies *λ ∈* [0, 1], thus 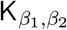 is row-stochastic. Because every row of R equals *r* and *r* sums to one, R^2^ = R. Therefore,

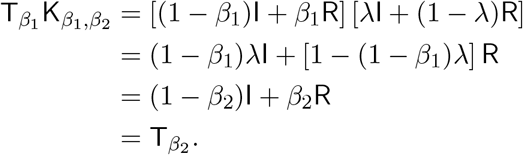

If *β*_1_ = 1, then *β*_2_ = 1, so the result is immediate.

#### Theorem 3.1

(Feasible-set contraction under garbling). Suppose that T_2_ = T_1_K for a row-stochastic post-processing kernel K, and suppose that the policy class *P* is closed under this post-processing. Then

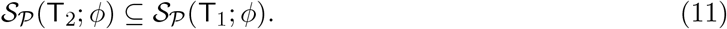

*Proof*. Consider any policy *π ∈ P* applied after observing the signal generated by T_2_. Under T_1_, the decision-maker can first generate a secondary signal through K and then apply *π*. Because T_2_ = T_1_K, the resulting joint distribution of the latent state, secondary signal, and action is identical to that obtained under T_2_. By post-processing closure, the composed policy belongs to *P* . Hence every expected utility pair attainable under T_2_ is also attainable under T_1_.

#### Corollary 3.1

(Monotonicity of the host-benefit frontier and MIDL). Under the conditions of Theorem 3.1,

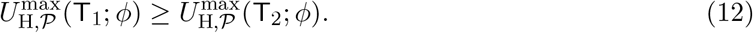

In particular, every measurement channel T is a garbling of the identity experiment because T = IT. Therefore,

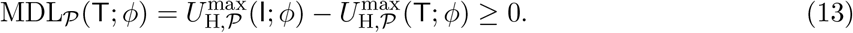

For the scalar family, if 0 *≤ β*_1_ *≤ β*_2_ *≤* 1, then

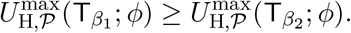

*Proof*. The feasible-set inclusion in Theorem 3.1 implies that maximizing the *U*_H_ coordinate over the smaller set cannot yield a larger value. The identity-channel and scalar-channel statements then follow from T = IT and Proposition 3.1, respectively.

Thus, MIDL is the decision loss attributable to using T in place of perfect observation. Under the stated closure conditions, improving information can reduce this loss without forcing the decision-maker to select a more intensive intervention. For comparison, *H*_*P*_ (T; *ϕ*) *−H*_*P*_ (R; *ϕ*) measures the benefit supplied by T relative to an uninformative channel. Along the scalar family, this delivered benefit and MIDL sum to the fixed quantity *H*_*P*_ (I; *ϕ*) *− H*_*P*_ (R; *ϕ*).

### 3.3 Scope of the Blackwell ordering

The general contraction theorem applies only when the policy class is closed under the relevant post-processing. The following result establishes closure for the convexified monotone class under the scalar affine kernels used above.

#### Proposition 3.2

(Closure of the convexified monotone class under scalar garbling). Let *ℳ* be the convex hull of nondecreasing deterministic policies. For a scalar affine post-processing kernel

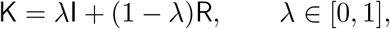

composition with K maps *ℳ* into itself.

*Proof*. It is sufficient to consider a deterministic nondecreasing policy *σ*. Its composition with K is

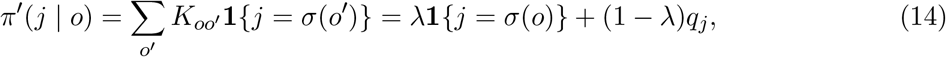

where 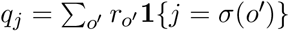 . The first term implements the original monotone policy *σ* and the second term is independent of the observed signal and can be implemented as an ex ante mixture of the constant policies *σ*_*j*_(*o*) *≡ j*, each of which is nondecreasing. Thus, the composed policy is a convex combination of nondecreasing deterministic policies and belongs to *ℳ*. Linearity extends the result to randomized mixtures in *ℳ*.

Consequently, the scalar feasible-set contraction and MIDL monotonicity results apply directly to the convexified monotone policy class used throughout the binary- and finite-state analyses.

## 4 A minimal binary-state model

### 4.1 Posterior policy regimes

Let *K* = 2, with 0 *< α <* 1, prior *P* (*X* = 1) = *α* and *P* (*X* = 0) = 1 *− α*. Use *r* = (1*/*2, 1*/*2), thus

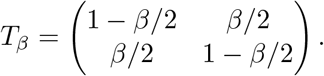

The marginal probability of the high observed signal is

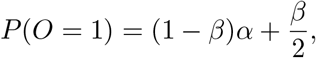

but the action after a single observation depends on the posterior state probability. Normalize the aggressive intervention’s gross benefit to one and its burden to *η ∈* (0, 1). Conditional on a signal *o*, the aggressive intervention has expected net host utility *P* (*X* = 1 | *O* = *o*) *− η*.

#### Lemma 4.1 (Posterior probabilities).

For *β ∈* [0, 1],

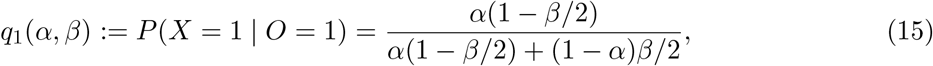

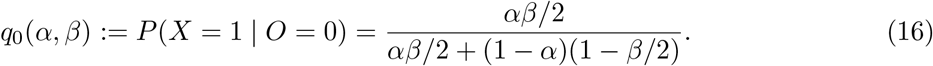

The aggressive tier is optimal after signal *o* if and only if *q*_*o*_(*α, β*) *≥ η*.

Define

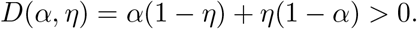

#### Proposition 4.1 (Optimal binary-state policy regimes).

For 0 *< α, η <* 1, define

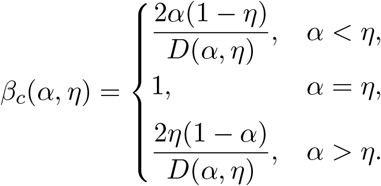

For 0 *≤ β < β*_*c*_(*α, η*), the host-benefit-maximizing monotone policy is observation-dependent: choose the conservative tier after *O* = 0 and the aggressive tier after *O* = 1.

If *α < η* and *β > β*_*c*_(*α, η*), the optimal policy is always conservative. If *α > η* and *β > β*_*c*_(*α, η*), the optimal policy is always aggressive. At *β* = *β*_*c*_(*α, η*), the observation-dependent policy ties the corresponding constant policy.

*Proof*. Because *q*_1_(*α, β*) *≥ q*_0_(*α, β*), only three monotone policies are possible. If *α < η*, then *q*_0_(*α, β*) *< η* for *β <* 1, and solving *q*_1_(*α, β*) = *η* gives

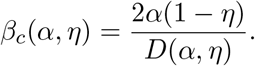

If *α > η*, then *q*_1_(*α, β*) *> η* for *β <* 1, and solving *q*_0_(*α, β*) = *η* gives

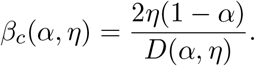

When *α* = *η*, the observation-dependent policy remains optimal until the signal becomes uninformative at *β* = 1.

#### Remark 4.1 (Symmetric prior).

When *α* = 1*/*2, *β*_*c*_(1*/*2, *η*) = 2 min *{η*, 1 *− η}* . For *η <* 1*/*2, the observation-dependent policy ties the always-aggressive policy at *β* = 2*η*. For *η >* 1*/*2, it ties the always-conservative policy at *β* = 2(1 *− η*). When *η* = 1*/*2, the observation-dependent policy remains optimal until complete information loss at *β* = 1.

### 4.2 Feasible-set geometry and measurement-induced decision loss

Denote the always-conservative, observation-dependent, and always-aggressive policies by *A, B*, and *C*, respectively. For the minimal binary-state case, specialize the general ecological utility to *ℓ*_0_ = *δ*_0_ = 0 and *ℓ*_1_ = *δ*_1_ = *d ≥* 0. This aligned normalization is consistent with the baseline condition *δ*_0_ = 0 in Assumption 2.1. Thus,

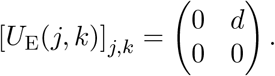

#### Proposition 4.2

(Binary-state feasible triangle and closed-form MIDL). The three outcome points in (*U*_E_, *U*_H_) coordinates are

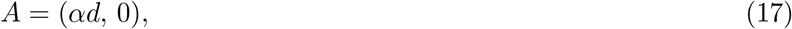

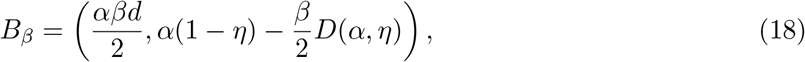

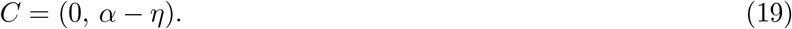

Hence, *S*_*ℳ*_(T_*β*_; *ϕ*) = conv*{A, B*_*β*_, *C}*, possibly degenerately. Moreover,

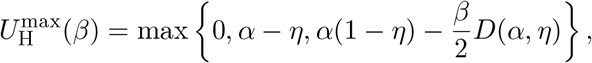

And

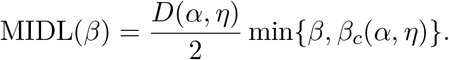

Thus, MIDL is nonnegative, nondecreasing, concave, and piecewise linear, with at most one interior kink. When *α* ≠ *η*, the kink at *β*_*c*_ *<* 1 is a switch from the observation-dependent optimum to a constant rule. When *α* = *η, β*_*c*_ = 1 and MIDL is linear on the entire interval; the policies tie only at its endpoint.

### Proof of Proposition 4.2 is provided in Section S1.2

Figure 1 illustrates how information loss changes feasible-set geometry in the minimal binary-state model (see Section S2.1). The constant-policy vertices correspond to decisions that do not depend on the observed signal: *A* is the always-conservative policy and *C* is the always-aggressive policy. Because these policies ignore the measurement, their outcomes are independent of *β*. The observation-dependent vertex *B*_*β*_, by contrast, represents the policy that selects the conservative tier after the low signal and the aggressive tier after the high signal. Under scalar blurring, *B*_*β*_ moves affinely toward the line segment joining the two constant-policy vertices, while *A* and *C* remain fixed. The resulting feasible triangle therefore contracts as the measurement becomes less informative.

**Figure 1.**
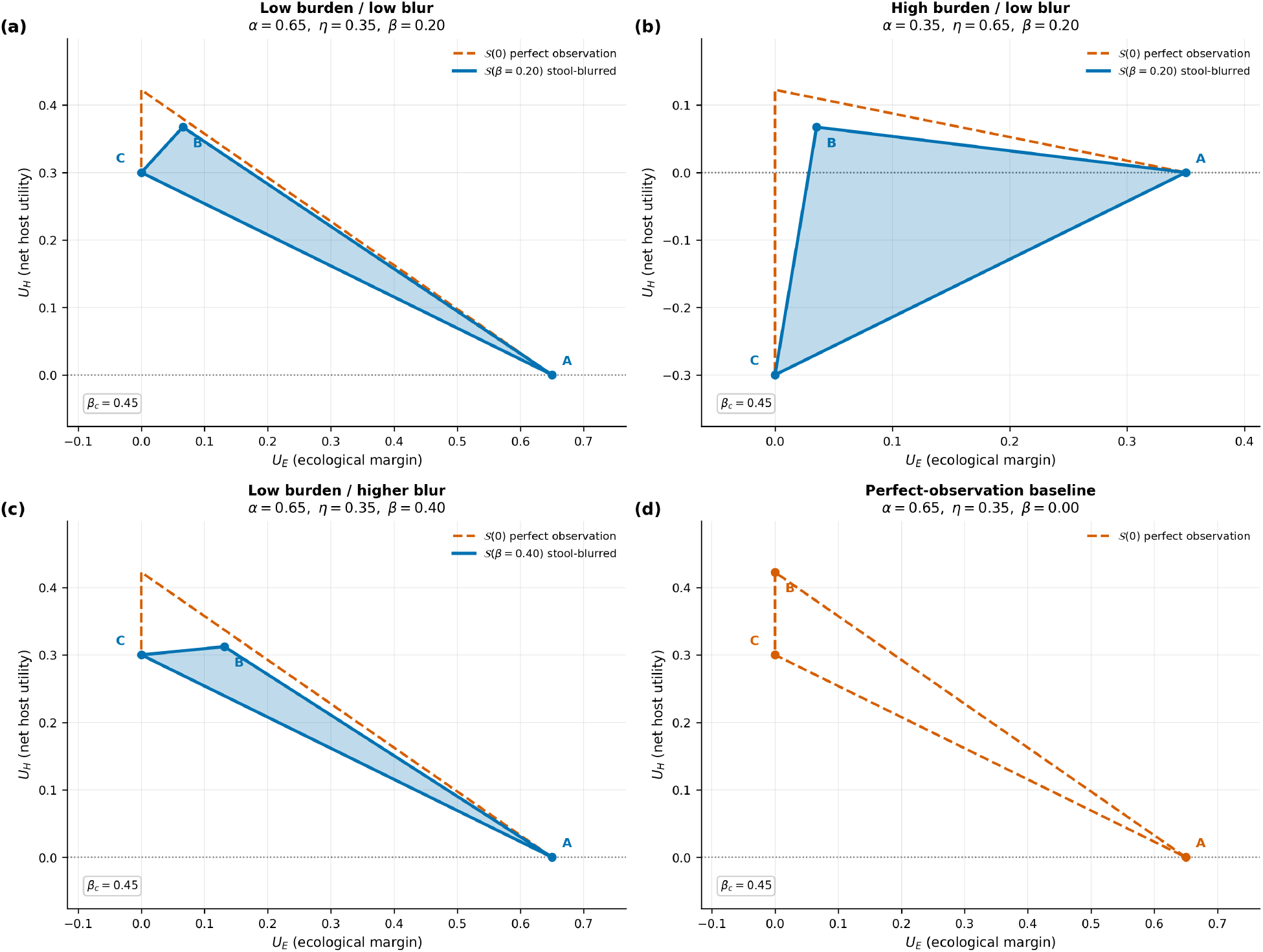
Feasible-set geometry in the binary-state model. The dashed outline shows the identity-channel feasible set, and the filled region shows the feasible set under information loss. Vertices *A, B*_*β*_, and *C* correspond to the always-conservative, observation-dependent, and always-aggressive policies. Panels vary the prior, intervention burden, and information-loss level as specified in Table S1.

The geometry depends jointly on the prior probability of the high latent state, *α*, and the normalized burden of the aggressive intervention, *η*. These parameters determine both the location of the constant-policy vertices and the critical information-loss level *β*_*c*_(*α, η*) at which the observation-dependent policy ties the relevant observation-independent policy. In the high-burden case with *α < η*, the always-aggressive vertex satisfies *U*_*H*_(*C*) = *α − η <* 0; the portion of the feasible set below *U*_*H*_ = 0 therefore represents attainable but host-disadvantageous policies and their randomized mixtures. At *β* = 0, the measurement channel is the identity and the feasible set equals the perfect-observation benchmark. As *β* increases, the attainable benefit of observation-dependent personalization declines until the optimal policy switches to a constant rule. This binary geometry anticipates the finite-state result that feasible sets are convex hulls of monotone-policy outcomes and that piecewise-linear changes in VoGM arise from the affine movement and switching of policy-specific vertices.

### Measurement-induced decision-loss landscape

Figure 2 illustrates how the closed-form MIDL derived in Proposition 4.2 depends on the prior high-state probability *α*, intervention burden *η*, and information-loss level *β*.

**Figure 2.**
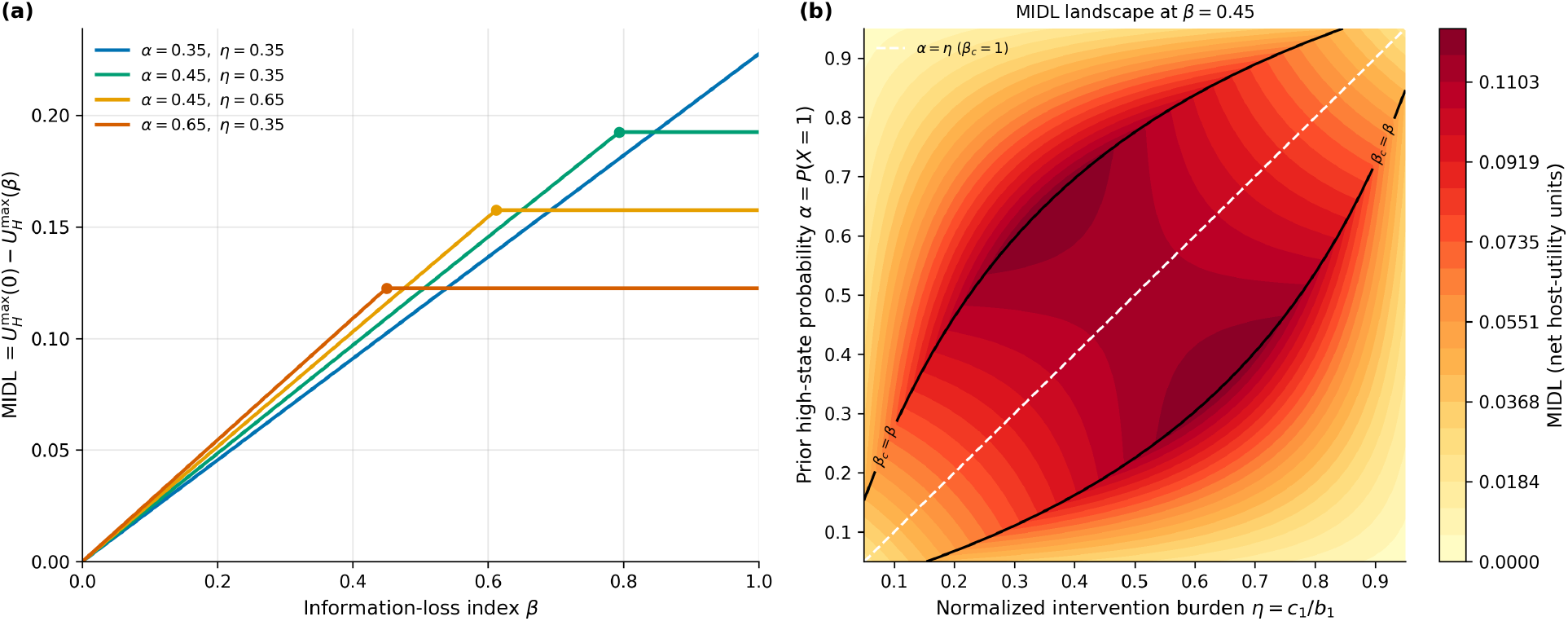
Measurement-induced decision loss in the binary-state model. **(a)** MIDL as a function of the scalar information-loss parameter *β* for selected values of the prior high-state probability *α* and normalized aggressive-intervention burden *η* = *c*_1_*/b*_1_. MIDL increases linearly while the observation-dependent policy maximizes net host benefit and plateaus after that policy ties the relevant constant policy at *β*_*c*_(*α, η*). Markers indicate interior policy-switch points; when *α* = *η*, the tie occurs only at *β*_*c*_ = 1.**(b)** MIDL over the (*η, α*) plane at *β* = 0.45. The black contour marks *β*_*c*_(*α, η*) = 0.45, separating parameter combinations for which the observation-dependent policy remains optimal from those for which the optimum has switched to a constant policy. The dashed diagonal denotes *α* = *η*, where *β*_*c*_ = 1.

**Figure 3.**
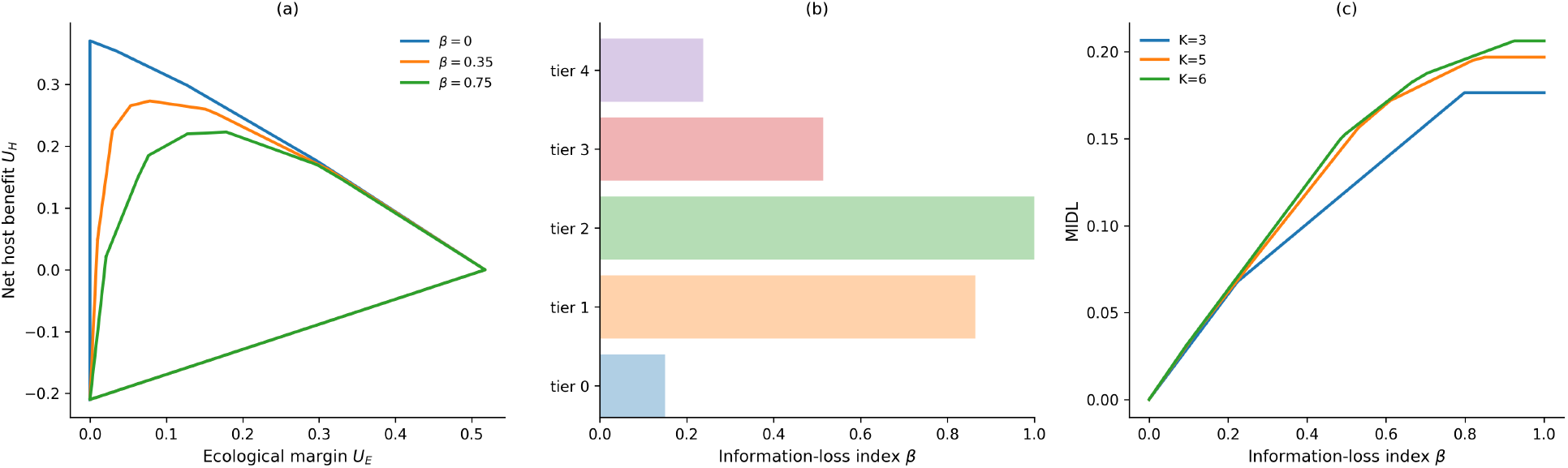
Finite-state feasible-set geometry and policy switching. **(a)** Feasible outcome sets for the *K* = 5 system under increasing scalar information loss, optimized over randomized mixtures of monotone policies. **(b)** Posterior-optimal intervention-tier viability using the same nonuniform prior and host-utility parameters as panel (a). A tier is viable when it belongs to the posterior-utility argmax for at least one observed signal; all tied maximizers are included. This diagnostic does not impose monotonicity across signals. **(c)** MIDL for *K* = 3, 5, 6 under the supplementary rank-based parameterizations. Kinks indicate switches between maximizing monotone policies. Larger state spaces can permit more switches but do not guarantee them.

For fixed (*α, η*), MIDL increases with slope *D*(*α, η*)*/*2 until *β* reaches *β*_*c*_(*α, η*). Beyond this point, the maximizing policy is observation-independent, so further information loss does not reduce the maximum attainable host benefit. At fixed *β*, MIDL varies across the (*α, η*) plane because the prior and intervention burden jointly determine the benefit of state-dependent intervention and the location of the policy switch. MIDL is small when the imperfect channel retains nearly all host benefit attainable under perfect observation, and large when observation loss prevents a materially better state-dependent policy. A plateau means that further degradation no longer changes the optimal host-benefit value for this decision problem; the channel may still contain information useful for other purposes.

## 5 *K*-state feasible-set theory

The convex-hull representation of randomized finite policies, the combinations-with-repetition count, and the upper-envelope argument below use standard finite-dimensional decision and convexity facts. Here they give explicit attainable-set and MIDL calculations for the selected ordinal utility structure. The number of policies bounds the potential complexity but does not guarantee a particular number of optimal-policy switches.

The binary-state model exposes the threshold geometry. The general finite-state model retains the same decision structure but contains more monotone policies and can therefore generate a more complex feasible outcome set.

Let

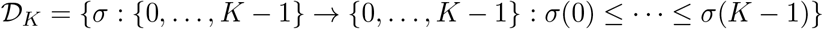

denote the class of deterministic nondecreasing policies. Each *σ ∈ D*_*K*_ induces the stochastic decision rule *π*_*σ*_(*j* | *o*) = **1***{j* = *σ*(*o*)*}*.

Let *ℳ*_*K*_ = conv *{π*_*σ*_ : *σ ∈ D*_*K*_*}* denote the class of randomized mixtures of these deterministic policies. Equivalently, every *π ∈ ℳ*_*K*_ can be written as

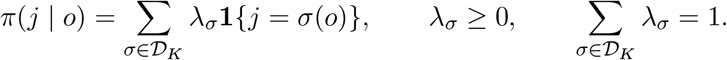

Such a rule can be implemented by drawing a deterministic monotone policy according to the probabilities *{λ*_*σ*_*}* before observing the measurement signal.

### Theorem 5.1

(Finite-state feasible-set representation). The number of deterministic nondecreasing policies is

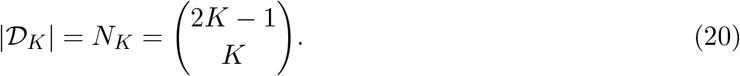

For fixed prior *ϕ*, measurement channel T, and utility parameters, define the outcome generated by *σ ∈ D*_*K*_ as

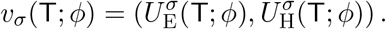

Then

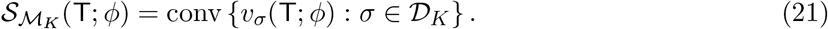

Consequently, the feasible outcome set is a compact convex polytope in ℝ^2^, generated by at most *N*_*K*_ distinct outcome points. Depending on the prior, channel, and utilities, it may be a polygon, line segment, or single point.

*Proof*. A deterministic nondecreasing policy is a weakly increasing sequence (*σ*(0), …, *σ*(*K* 1)) of length *K*, with entries in *{* 0, …, *K −* 1*}* . The number of such sequences equals the number of multisets of size *K* drawn from *K* possible values. By the standard combinations-with-repetition formula,

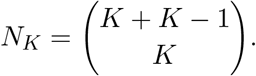

Each deterministic policy *σ* generates one expected utility pair *v*_*σ*_(T; *ϕ*). If a policy *π ∈ ℳ*_*K*_ uses mixture weights *{λ*_*σ*_*}*, linearity of expected utility gives

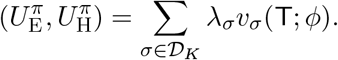

Thus, every randomized mixture generates a convex combination of the deterministic-policy outcomes. Conversely, every such convex combination can be implemented by drawing a deterministic policy according to *{ λ*_*σ*_} before observing the signal. Therefore, the feasible set is exactly the convex hull in Eq. (21).

The quantity *N*_*K*_ counts the candidate deterministic policies and therefore bounds the number of policy-generated outcome points. It does not generally equal the number of vertices of the feasible set: distinct policies may generate identical outcome points, and some policy-generated points may lie strictly inside the convex hull.

### Theorem 5.2

(Piecewise-linear finite-state frontier). Under the scalar affine channel, Eq. (7), the outcome generated by every fixed *σ ∈ D*_*K*_ is affine in *β*. Consequently,

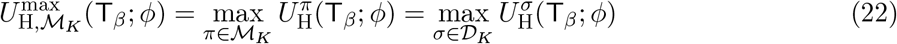

is convex and piecewise linear in *β*. Therefore,

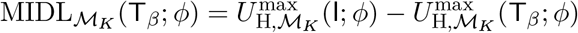

is concave and piecewise linear in *β*.

*Proof*. For a fixed deterministic policy *σ*, let

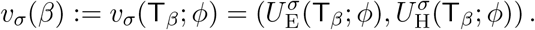

By linearity of expected utility in the measurement channel Eq. (7) gives

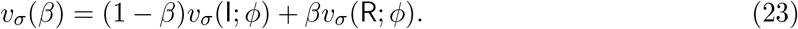

Therefore, both coordinates of *v*_*σ*_(*β*) (the expected ecological margin and the expected net host benefit) are affine functions of *β*.

Because expected host benefit is linear in the policy mixture, maximizing over *ℳ* _*K*_ is equivalent to maximizing over its deterministic generating policies *D* _*K*_. Since *D* _*K*_ contains only *N*_*K*_ policies, the host-benefit frontier is the maximum of finitely many affine functions of *β*. It is therefore convex and piecewise linear, with changes in slope occurring when the maximizing deterministic policy changes. Subtracting this frontier from the constant perfect-observation benchmark produces a concave, piecewise-linear MIDL.

Figure **??** illustrates the finite-state results in synthetic ordered-state systems (see Section S2.2). Panels (a) and (b) share the same *K* = 5 benefit vector, burden vector, reference channel, and nonuniform prior. Panel (a) displays contraction of the convexified monotone-policy feasible set. Panel (b) identifies tiers that maximize posterior expected host benefit for at least one signal; this is a signalwise Bayes diagnostic without a monotonicity restriction on the collection of maximizing actions. Consequently, posterior tier viability need not imply use of that tier by a globally optimal monotone policy. Panel (c) compares *K* = 3, 5, 6 systems using its stated rank-based prior rule and illustrates changes in MIDL slope when a maximizing monotone policy switches.

The figure shows that increasing the number of states changes the combinatorial complexity of the decision problem but not its basic geometry. Each deterministic monotone policy generates an outcome trajectory that is affine in *β*, and the feasible set is the convex hull of these policy-specific outcomes. The optimal host-benefit frontier is therefore the upper envelope of finitely many affine functions. Larger state spaces can support more feasible vertices and more policy switches, but feasible-set contraction under scalar garbling and piecewise-linearity of MIDL remain unchanged.

## 6 State-specific information loss

The scalar channel assigns the same information-loss level to every latent state (see Eq. (7)). In practice, measurement fidelity may depend on the underlying functional configuration because spatial localization, adhesion, growth, and transit effects differ across communities. We therefore allow the information-loss parameter to vary by latent state; we replace the common scalar *β* with the vector ***β***.

### Definition 6.1

(State-specific affine channel). Let ***β*** = (*β*_0_, …, *β*_*K−* 1_) *∈* [0, 1]^*K*^ and let *r ∈*Δ^*K−*1^ be the reference signal distribution. The state-specific affine measurement channel is

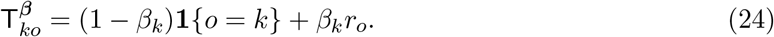

Thus, conditional on latent state *k*, the observed signal equals the state-matched category with weight 1 *− β*_*k*_ and is replaced by a draw from the reference distribution with weight *β*_*k*_.

Each row of T^***β***^ is a convex combination of the point mass *e*_*k*_ and *r*, so the channel is row-stochastic. The scalar channel is recovered when all *β*_*k*_ are equal, and the identity channel is recovered when all *β*_*k*_ = 0.

Because T^***β***^ = IT^***β***^, every state-specific channel is a garbling of the identity experiment. Therefore, under the closure condition of Theorem 3.1, *S*_*P*_ (T^***β***^ ; *ϕ*) *⊆ S*_*P*_ (I; *ϕ*), and VoGM relative to perfect observation is nonnegative.

### Remark 6.1

(No automatic ordering among state-specific channels). Unlike the scalar family

*{*T_*β*_*}*_*β∈*[0,1]_, the state-specific family is not generally totally ordered by Blackwell informativeness. In particular, even if 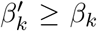 for every *k*, it does not generally follow that 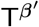 is a garbling of T^***β***^ . Componentwise larger information-loss parameters therefore do not, by themselves, guarantee nesting of the corresponding feasible sets. Comparisons between two state-specific channels require either an explicit stochastic kernel K satisfying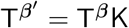, or direct evaluation of their attainable feasible sets and decision frontiers. An explicit counterexample is given in Proposition S1.1.

## 7 Multi-segment stool aggregation

The categorical decision model developed above represents the intervention-relevant condition by a latent state *X ∈{*0, …, *K−* 1*}*, with population prior *ϕ*. Stool, however, can also be modeled directly at the compositional level as an aggregate of material contributed by multiple intestinal segments. To distinguish these two levels of description, let

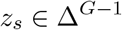

denote the functional-guild composition of intestinal segment *s*, where *G* is the number of biological guilds. Thus, *z*_*s*_ is a within-segment composition, whereas *ϕ* remains a prior distribution over categorical latent states.

Suppose that segment *s* contributes weight *w*_*s*_ to the stool aggregate and is represented with segment-specific information-loss level *β*_*s*_. A common reference profile *r ∈* Δ^*G−*1^ represents the limiting composition toward which segment-specific observations are blurred. The following algebraic identity characterizes compositional approximation error when this heterogeneous aggregation process is replaced by a single blur applied to the weighted mean composition. It does not itself define *P* (*O* |*X*) or determine MIDL. A known noiseless affine map *m* = (1 *− β*)*z* + *βr*^comp^ is invertible for *β <* 1; a decision-level information-loss interpretation therefore additionally requires a probabilistic observation mechanism or an explicit categorization of the observed profile. Connecting the aggregation model to that categorical experiment requires calibration beyond the identity established here.

### Proposition 7.1

(Multi-segment blur–composition covariance identity). Let *s* = 1, …, *S* index intestinal segments, with

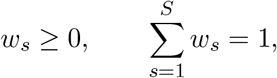

segment compositions *z*_*s*_ *∈* Δ^*G−*1^, segment-specific information-loss levels *β*_*s*_ *∈* [0, 1], and a common reference profile *r*^comp^ *∈* Δ^*G−*1^. Define

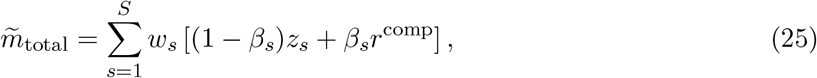

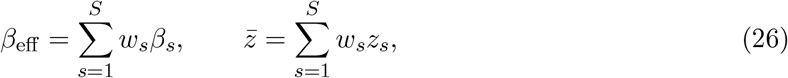

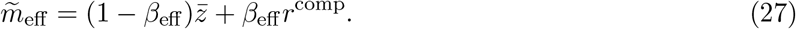

Then

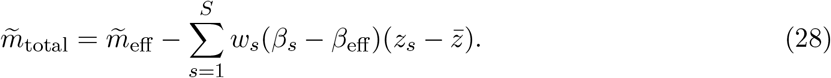

The correction term in Eq. (28) is a vector-valued weighted covariance between segment-specific information loss and segment-specific composition. Its magnitude and direction therefore depend on how deviations in *β*_*s*_ align with deviations in *z*_*s*_, not on the variance of the blur levels alone.

### Corollary 7.1

(Exactness of the single-blur representation). The heterogeneous aggregate equals the mean-level single-blur representation, 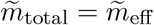, if and only if

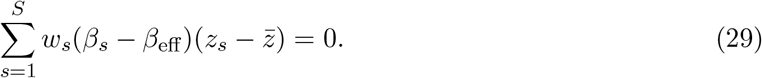

In particular, the equality holds if either all segment compositions are equal or all segment-specific information-loss levels are equal.

### Corollary 7.2

(Norm bound). For any norm on ℝ^*G*^,

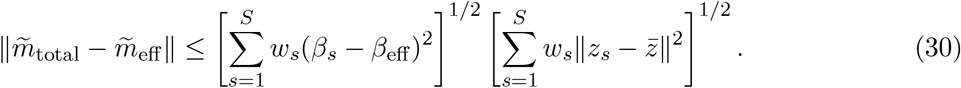

Proofs of Proposition 7.1 and Corollaries 7.1 and 7.2 are provided in Section S1.3.

Figure 4 illustrates the identity in a deterministic synthetic system with four intestinal segments and four functional guilds (see Section S2.3 for the details). Variation in segment-specific information loss alone does not produce an approximation error: when all segments have the same composition, the covariance term vanishes even though the *β*_*s*_ values differ. As segment compositions become increasingly heterogeneous, their alignment with segment-specific blur produces a nonzero discrepancy between the heterogeneous aggregate and the single-blur representation. The exact discrepancy remains below the bound in Eq. (30).

**Figure 4.**
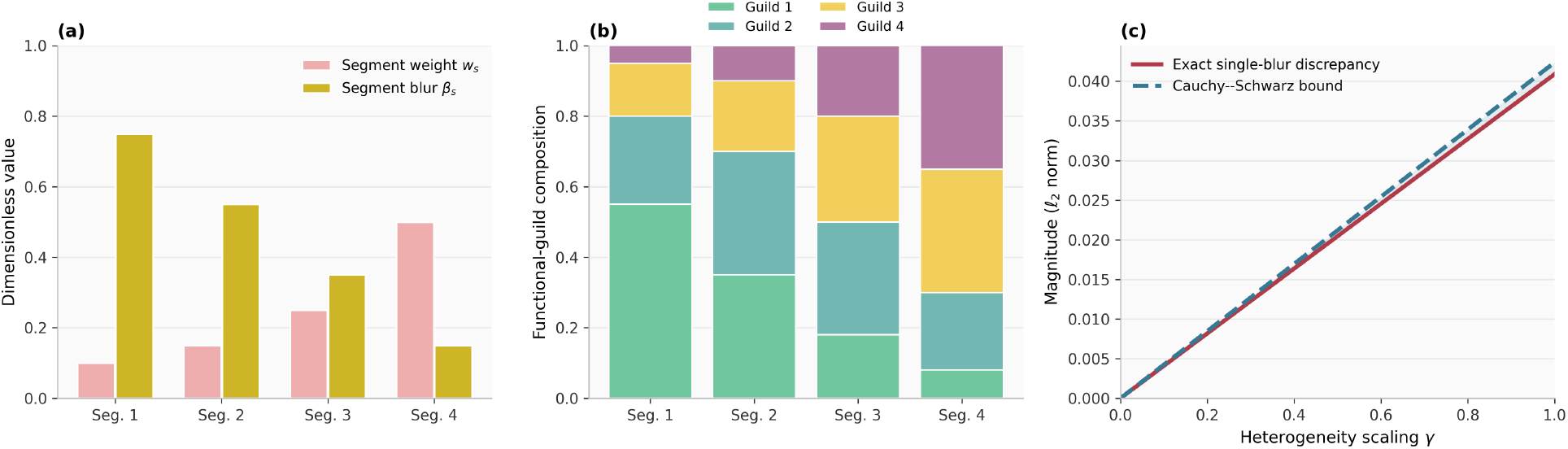
Multi-segment blur–composition covariance identity. Synthetic illustration with *S* = 4 intestinal segments and *G* = 4 functional guilds. **(a)** Segment contribution weights *w*_*s*_ and segment-specific information-loss levels *β*_*s*_. Their weighted mean is *β*_eff_ = Σ_*s*_ *w*_*s*_*β*_*s*_ = 0.32. **(b)** Segment-specific functional-guild compositions *z*_*s*_, chosen to represent a stylized proximal–distal upper bound in (30) as the heterogeneity-scaling parameter interpolates from identical segment compositional gradient. **(c)** Exact Euclidean discrepancy 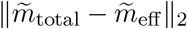 and the Cauchy–Schwarz compositions (*γ* = 0) to the full synthetic gradient (*γ* = 1). The discrepancy is zero at *γ* = 0, despite heterogeneous blur levels, because the blur–composition covariance term vanishes. Both curves are linear in this illustration because the segment compositions are varied through an affine interpolation in *γ*. The parameterization is illustrative and is not fitted to empirical intestinal-segment data.

## 8 Ecological-margin constraints

### 8.1 Constraints on attainable outcomes

The feasible set records expected ecological margin and net host benefit as separate outcome coordinates. In some applications, comparing these coordinates is insufficient: an intervention may provide high expected host benefit while failing to preserve a minimum acceptable ecological margin. This motivates restricting the feasible set to policies satisfying an explicit ecological-margin floor.

#### Definition 8.1

(Ecologically admissible feasible set). For a policy class *P*, prior *ϕ*, measurement channel T, and ecological-margin floor *τ*, define

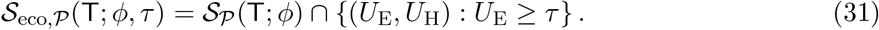

#### Proposition 8.1

(Convexity under an ecological-margin floor). If *S* _*P*_ (T; *ϕ*) is convex, then _eco,_ *S*_*P*_ *S*_*eco,P*_ (T; *ϕ, τ* ) is convex. If *S*_*P*_ (T; *ϕ*) is compact, then the ecologically admissible set is compact, although it may be empty.

*Proof*. The set *{* (*U*_E_, *U*_H_) : *U*_E_ *≥ τ}* is a closed convex half-space. Its intersection with a convex set is convex, and its intersection with a compact set is compact.

#### Definition 8.2

(Expected-margin-constrained frontier). Whenever *S*_*P*_ (T; *ϕ*) is compact and *S*_eco,*P*_ (T; *ϕ, τ* ) is nonempty, define

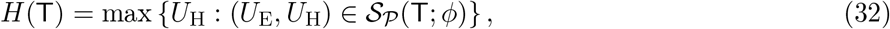

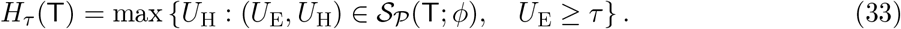

The ecological-screening gap is

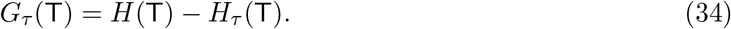

#### Proposition 8.2

(Information and the ecological-screening gap). Suppose that T_2_ is a garbling of T_1_ and that *P* is closed under the corresponding post-processing. Then *H*(T_1_) *≥ H*(T_2_). Whenever both ecologically admissible sets are nonempty, *H*_*τ*_ (T_1_) *≥H*_*τ*_ (T_2_). However, feasible-set inclusion alone does not determine the ordering of *G*_*τ*_ .

*Proof*. The garbling relation and post-processing closure imply *S*_*P*_ (T_2_; *ϕ*) *⊆ S*_*P*_ (T_1_; *ϕ*). Maximizing *U*_H_ over these nested sets gives the first inequality. Intersecting both sets with the same half-space *U*_E_ *≥ τ* preserves inclusion, which gives the second inequality.

To see why the difference between the two frontiers is not ordered by set inclusion alone, consider the nested compact convex sets *S*_2_ = *{* (1, 1) *}* and *S*_1_ = conv *{* (1, 1), (1, 1.1), (0, 2) *}*, and set *τ* = 1*/*2. For _2_, *H*_2_ = *H*_*τ*,2_ = 1, so the gap is zero. For _1_, the unconstrained frontier is *H*_1_ = 2. The segment joining (1, 1.1) and (0, 2) intersects *U*_E_ = 1*/*2 at (1*/*2, 1.55), giving *H*_*τ*,1_ = 1.55 and *G*_*τ*,1_ = 0.45. Thus, both frontiers can improve while their difference increases. The binary model also supplies both directions of gap change under the same scalar ordering. For *α* = 0.60, *η* = 0.30, *d* = 1, and *τ* = 0.228, the plotted range 0 *≤ β≤* 0.55 has *U*_E_(*B*_*β*_) = 0.30*β < τ* . The constrained optimum lies on the segment joining *A* and *B*_*β*_, and hence

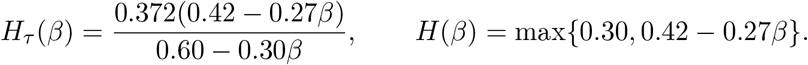

These formulas give *G*_*τ*_ (0) = 0.1596, *G*_*τ*_ (4*/*9) *≈* 0.06086, and *G*_*τ*_ (0.55) *≈* 0.06782. Thus, the gap decreases and then increases as information loss grows, although both frontiers are nonincreasing.

No monotonic ordering of the gap follows from feasible-set inclusion alone.

#### Remark 8.1

(Expected versus statewise ecological constraints). The floor in Eq. (31) constrains expected ecological margin. It may therefore admit a randomized or heterogeneous policy that combines low-margin and high-margin outcomes. If ecological protection must hold for each treated state, uniformly over uncertainty, or with a prescribed probability, the expected floor should instead be replaced by a statewise, robust, or chance constraint [Ben-Tal et al., 2009, Nemirovski and Shapiro, 2006]. These alternatives generally define different admissible sets.

### 8.2 Ecological reserve under community dynamics

To motivate why an ecological-margin floor may be desirable, consider the two-guild generalized Lotka–Volterra system [Stein et al., 2013, Venturelli et al., 2018]:

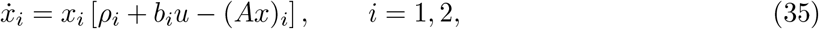

where *u* is intervention intensity, *ρ* is the baseline growth vector, *b* describes the intervention-induced growth shift, and *A* is the interaction matrix. An interior equilibrium satisfies

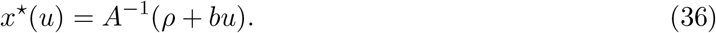

The condition *x*^*⋆*^(*u*) *>* 0 establishes coexistence feasibility but does not, by itself, establish local asymptotic stability. This distinction is important in microbiome ecology, where community persistence, interaction structure, and dynamical stability are related but non-equivalent properties [Coyte et al., 2015].

For an arbitrary equilibrium *x*^*⋆*^, including a boundary equilibrium, the Jacobian is

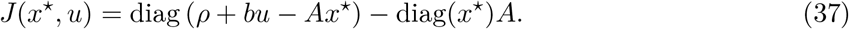

At a positive interior equilibrium, the first term vanishes, so local asymptotic stability requires

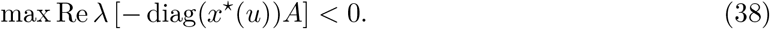

For a boundary equilibrium, stability must instead be evaluated using the full Jacobian in (37), because the invasion growth rate of an absent guild need not vanish.

Figure 5 separates the biological motivation for ecological screening from its decision-theoretic implementation (see Section S2.4 for the details). Panel (a) shows a genuine change in ecological regime, from stable coexistence to a stable boundary state. Panels (b)–(c) represent such concerns through an expected ecological-margin floor using the abstract ecological-margin utility.

**Figure 5.**
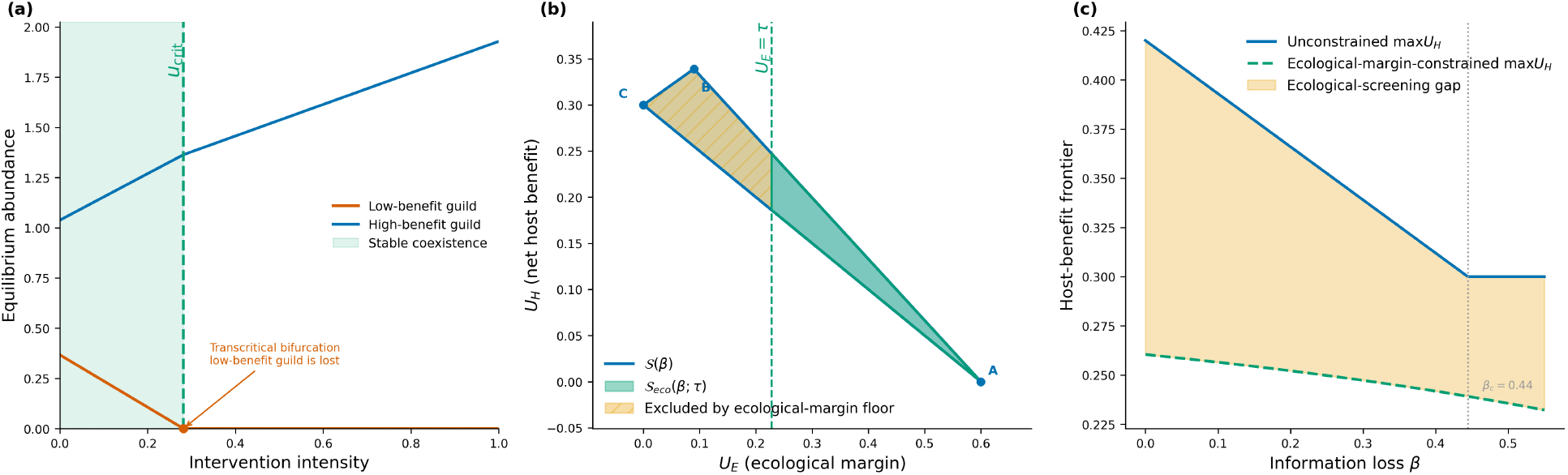
Ecological-margin screening gap. **(a)** Locally stable equilibria of a two-guild generalized Lotka–Volterra model as intervention intensity *u* increases. The shaded region denotes stable positive coexistence; at *u*_crit_, Guild 1 is lost and the interior and boundary branches exchange stability through a transcritical bifurcation. Guild labels do not imply a host-benefit ranking. **(b)** Binary-state feasible set at fixed information loss, intersected with the expected ecological-margin constraint *U*_E_ *≥ τ* . The green region is admissible, whereas the hatched region is excluded by the constraint. **(c)** Unconstrained and ecological-margin-constrained maximum net host benefit as information loss *β* varies. The shaded difference is the ecological-screening gap, and the dotted line marks the host-benefit policy-switch point *β*_*c*_.

The floor may remove part of the attainable feasible set and can therefore lower the maximum admissible host benefit. Because the ecological utility is truncated at zero, it records remaining modeled reserve; its expected-value floor does not guarantee protection in every state. Under scalar garbling, more informative measurement weakly improves both the unconstrained and constrained frontiers, but the ecological-screening gap between them need not vary monotonically.

## 9 Discussion

The decision-theoretic insight is that a measurement is useful for intervention when it preserves the distinctions that change preferred actions and attainable outcomes. Faithful reconstruction of the entire gut ecosystem is neither necessary nor sufficient to establish that usefulness for a specified decision problem. MIDL makes the consequence of imperfect observation explicit: it is the reduction in maximum attainable expected net host benefit relative to perfect observation. A larger MIDL indicates greater forgone benefit, and a smaller MIDL indicates limited loss only for the specified population, interventions, utilities, and policy class. It does not establish measurement accuracy or adequacy for other scientific tasks.

The feasible set provides information beyond the host-benefit maximum. Two measurement channels may have similar MIDL values while supporting different trade-offs between net host benefit and ecological margin. The same feasible set may also lead to different selected decision rules when greater weight is placed on treatment burden, ecological protection, or patient preference. By separating attainable outcomes from preferences over those outcomes, the framework permits measurement strategies to be compared without first reducing the decision problem to a single scalar objective.

Under scalar garbling and post-processing closure, a more informative measurement weakly expands the feasible outcome set and cannot reduce the maximum attainable host benefit [Blackwell, 1953]. This is an opportunity result: additional information enlarges the set of implementable decision rules but does not require selection of a more aggressive or ecologically unfavorable intervention. When an ecological-margin floor is imposed, the margin-constrained host-benefit frontier also weakly improves with information, although the difference between the constrained and unconstrained frontiers need not vary monotonically because information may expand high-benefit and ecologically admissible regions by different amounts.

The piecewise-linear structure of MIDL connects observation loss to decision-rule switching. While the same monotone policy maximizes host benefit, its value is affine in *β* and MIDL is linear. A change in slope marks a switch between optimal policy-value branches; the optimal value remains continuous even when the selected rule changes. In the binary model with *α* ≠ *η*, the switch is from a signal-dependent rule to a constant rule. The equal case *α* = *η* has no interior switch. Larger state spaces permit more candidate monotone mappings and can support additional switches, but their occurrence and spacing depend on the prior, utilities, and channel. In applications, proximity to a switch point and the stability of the selected action may therefore matter as much as average predictive performance.

### Biological interpretation

Translating these results into biological applications requires careful interpretation of both the latent state and the measurement channel. Real microbial communities contain multiple taxa and overlapping functional groups that vary continuously in abundance and behavior. The finite-state representation is therefore a decision-oriented abstraction. The latent state retains only the features needed for the modeled intervention and may represent metabolic capacity, intervention response, ecological risk, or another clinically meaningful quantity.

The measurement channel summarizes how each latent intervention-relevant state generates a distribution of observed signals. Rather than modeling every taxon, metabolite, and spatial process explicitly, it represents their combined effect as a probabilistic relationship between the underlying state and the stool-derived observation. This abstraction is motivated by the spatial heterogeneity of the gut microbiome [Donaldson et al., 2016, Shalon et al., 2023, Folz et al., 2023]. Its purpose is to determine whether stool preserves the distinctions among latent states that warrant different interventions.

The scalar affine channel provides an analytically transparent benchmark because it represents a single direction of increasing information loss and induces a total Blackwell order. Biological measurement processes, however, may be state-specific. Some functional states may be preferentially retained or obscured because their organisms or metabolites differ in localization, adhesion, growth, shedding, degradation, oxygen tolerance, or transit sensitivity. The state-specific extension represents this heterogeneity and shows that increasing the loss assigned to each state does not necessarily produce a uniformly less informative measurement. State-specific channels must therefore be compared through an explicit garbling relation or by directly evaluating the decisions and feasible outcomes they support.

Measurement heterogeneity also arises from aggregation across compositionally distinct gut regions. The multi-segment result shows that applying one effective blur to the weighted mean composition reproduces the heterogeneous aggregate only when the weighted covariance between segment-specific information loss and segment composition is zero. Variation in regional recovery alone is therefore insufficient to determine aggregation error; the error arises when regions with different compositions are represented with systematically different fidelity in stool. Paired intestinal– stool studies should consequently characterize regional contributions, compositions, and measurement processes jointly.

These results suggest that further sampling should be prioritized when it can distinguish latent states associated with different preferred interventions or clarify whether candidate interventions satisfy an ecological constraint. Regional gut sampling, repeated stool collection, multi-omic assays, and combinations of measurement modalities can then be compared through their effects on MIDL, the feasible outcome set, and the ecologically admissible frontier.

Finally, the ecological-margin coordinate requires careful interpretation. *U*_E_ is an abstract measure of ecological reserve remaining after intervention; it is not itself a direct estimate of resistance, resilience, return time, or local dynamical stability [Shade et al., 2012, Relman, 2012]. The gLV illustration motivates ecological screening by showing that increasing intervention intensity can alter the qualitative ecological regime, but it does not calibrate *U*_E_. In an empirical application, the ecological margin could instead be linked to a validated stability measure, the probability of maintaining a desired function, distance from an undesirable state, or a robust lower bound under model uncertainty.

### Future work

The results presented above are conditional on the specified state space, prior, measurement channel, and utility model. Applying the framework to a particular assay or intervention therefore requires empirical calibration of these components. Calibration involves defining intervention-relevant states, estimating how those states generate observed signals, and assigning host-benefit, burden, and ecological utilities. These quantities may be informed by mechanistic or clinically annotated response models, paired intestinal–stool observations, causal treatment effects, patient preferences, and expert elicitation. Repeated sampling can help distinguish measurement error from temporal variation, while uncertainty in the calibrated inputs should be propagated to MIDL, feasible frontiers, and decision-rule comparisons.

In this work, the scalar affine channel provides an analytically tractable benchmark, but more general state-dependent, subject-specific, or nonlinear channels can be evaluated within the same feasible-set framework. Their information ordering, however, must be established directly. Similarly, compositional transformations may support statistical analysis [Aitchison, 1986, Gloor et al., 2017], but they do not by themselves define stochastic measurement channels; Blackwell comparisons require an explicit probabilistic experiment that accounts for zeros and detection limits.

The expected ecological-margin floor is one possible safety criterion. Statewise, robust, or chance constraints may be preferable when protection must hold for each relevant state, under model uncertainty, or with a bounded failure probability. The framework could also be extended to sequential settings involving repeated measurement, Bayesian updating, adaptive intervention, and ecological monitoring. Candidate measurement strategies could then be compared by their effects on MIDL, the ecologically admissible frontier, or decision regret subject to sampling and resource constraints.

## 10 Conclusion

We developed a feasible-set theory for microbiome intervention under observation loss. The framework distinguishes the underlying functional-guild composition from a categorical, intervention-relevant latent state; represents stool measurement as a stochastic channel; and evaluates policies through net host benefit and ecological margin. Scalar affine information loss produces a Blackwell ordering and, under post-processing closure, nested feasible sets. The binary-state model yields closed-form policy switches and MIDL, while finite policy classes produce convex piecewise-linear host-benefit frontiers and concave piecewise-linear MIDL. State-specific loss need not be totally ordered, multi-segment aggregation introduces a blur–composition covariance correction, and ecological-margin floors restrict the set of admissible interventions. These results shift the evaluation of stool from the question of whether it is an imperfect proxy to the decision-relevant question of whether its information loss changes the interventions and trade-offs that remain attainable.

## Supplementary Information

### S1 Supplementary theoretical results

#### S1.1 State-specific information loss and Blackwell ordering

For the state-specific measurement channel, the probability of observing state *o* conditional on latent state *k* is

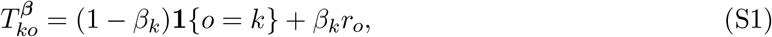

where *r* is the reference signal distribution and *β*_*k*_ *∈* [0, 1] is the state-specific information-loss parameter. Componentwise increases in ***β*** do not, in general, induce a Blackwell ordering of the resulting channels.

Proposition S1.1

(Componentwise blur does not imply Blackwell ordering). Let *K* = 3, *r* = (1*/*3, 1*/*3, 1*/*3), and consider

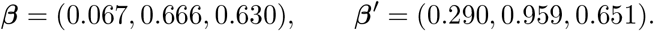

Although 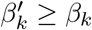 for every *k*, 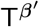 is not a garbling of T^***β***^ .

*Proof*. The matrix T^***β***^ is invertible. Therefore, any post-processing kernel K satisfying

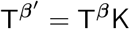

would necessarily equal

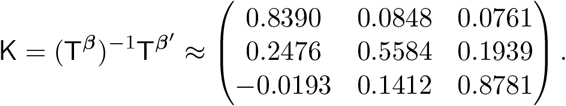

The rows sum to one, but the matrix contains a negative entry and is therefore not a stochastic kernel. Hence, 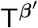 is not a garbling of T^***β***^ . The reverse candidate kernel 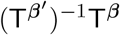 also contains a negative entry (approximately 0.5928), so neither direction defines a Blackwell garbling for this pair.

##### S1.2 Proof of feasible-set geometry for K=2

*Proof of* **??**. Under this normalization, the only potentially positive ecological margin is

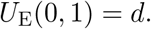

The always-conservative policy therefore has expected ecological margin

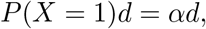

whereas the always-aggressive policy has expected ecological margin zero. Under the observation-dependent policy, the conservative tier is selected after *O* = 0. A positive ecological margin is therefore obtained only when *X* = 1 and *O* = 0, whose probability is

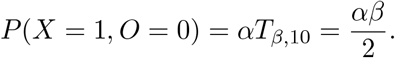

Hence its expected ecological margin is *αβd/*2.

The host-benefit coordinates follow by direct summation over

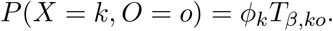

The perfect-observation policy *B*_0_ has host utility *α*(1 *− η*), which exceeds both constant-policy values for 0 *< α, η <* 1. The observation-dependent host utility decreases with slope *− D* (*α, η*)*/*2 until it meets the better constant policy at *β*_*c*_, after which additional information loss has no effect on the host-benefit optimum.

### S1.3 Proofs for the multi-segment aggregation results

***Proof of Proposition 7.1***. Expanding the heterogeneous aggregate gives

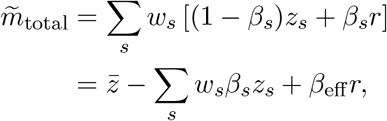

where 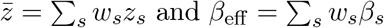 and *β*_eff_ = Σ_*s*_ *w*_*s*_*β*_*s*_. The weighted blur–composition covariance term satisfies

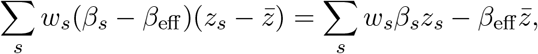

Because Σ_*s*_ *w*_*s*_ = 1. Therefore,

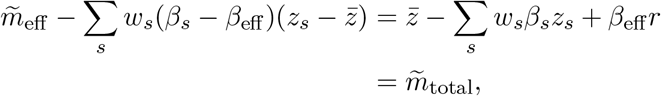

which proves the identity in Eq. (28).

*Proof of Corollary 7.1*

The result follows directly from Eq. (28). If all segment compositions are identical, then 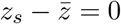 for every *s*. If all segments have the same information-loss level, then *β*_*s*_ *− β*_eff_ = 0 for every *s*. In either case, the covariance correction vanishes and the single-blur representation is exact.

*Proof of Corollary 7.2*

Let *a*_*s*_ = *β*_*s*_ *− β*_eff_ and 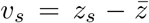. By Eq. (28) and the triangle inequality,

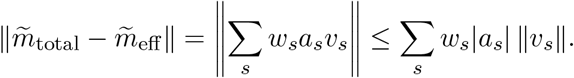

Applying the weighted Cauchy–Schwarz inequality yields

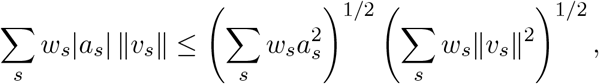

which proves Eq. (30).

### S2 Construction of the theoretical illustrations

All illustrations were generated from deterministic synthetic parameterizations chosen to demonstrate the theoretical behavior of the model. Unless explicitly stated otherwise, no empirical microbiome data were used for parameter fitting.

#### S2.1 Binary-state feasible-set illustration

The binary feasible-set illustration uses the parameterizations in Table S1, the ecological normalization *ℓ* = *δ* = (0, 1) (so *d* = 1), and the reference distribution *r* = (1*/*2, 1*/*2). The measurement channel is

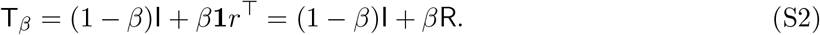

**Table S1.** Parameterization of the binary feasible-set illustration. Information loss is reported both directly and as a fraction of the policy-switch point *β*_*c*_(*α, η*).

| Panel | $\alpha$ | $\eta$ | $\beta/\beta_c$ | $\beta$ |
| --- | --- | --- | --- | --- |
| (a) Low burden, low blur | 0.65 | 0.35 | 0.45 | 0.2023 |
| (b) High burden, low blur | 0.35 | 0.65 | 0.45 | 0.2023 |
| (c) Low burden, higher blur | 0.65 | 0.35 | 0.90 | 0.4046 |
| (d) Identity-channel benchmark | 0.65 | 0.35 | 0 | 0 |

For each parameter setting, we evaluated the three deterministic monotone policies *σ*_*A*_ = (0, 0), *σ*_*B*_ = (0, 1) and *σ*_*C*_ = (1, 1) which correspond to always selecting the conservative tier, selecting the tier indicated by the observed signal, and always selecting the aggressive signal, and always selecting the aggressive outcomes were calculated from *P*(*X* = *k, O* = *o*) = *ϕ*_*k*_*T*_*β,ko*_ and *ϕ* = (1− *α, α*). The associated outcome points 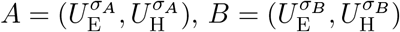, and 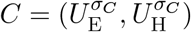 define

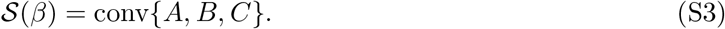

Figure **??** compares the feasible set under information loss with the identity-channel benchmark *S* (0). The panel parameterizations are reported in Table S1. Information loss is reported directly and as a fraction of the optimal policy-switch point *β*_*c*_(*α, η*) defined in Proposition 4.1.

#### S2.2 Finite-state illustrations

Figure **??** uses synthetic ordinal latent-state systems; the state count *K* is distinct from the number of biological guilds *G*. Panels (a) and (b) share the *K* = 5 parameterization

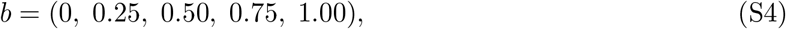

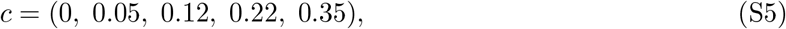

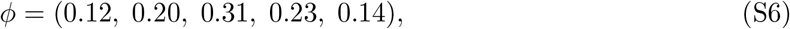

with a uniform reference distribution *r*_*k*_ = 1*/K* and ecological parameters *ℓ*_*k*_ = *δ*_*k*_ = *b*_*k*_. For each *β*, we enumerated all nondecreasing policies *σ* : *{*0, …, *K−* 1*} → {*0, …, *K –* 1}, calculated their expected (*U*_E_, *U*_H_) outcomes, and plotted the convex hull.

Figure **??**b uses exactly the same *b, c, ϕ*, and reference distribution as panel (a), including the nonuniform prior *ϕ* = (0.12, 0.20, 0.31, 0.23, 0.14). For every signal with positive marginal probability, the code calculates 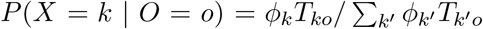 and the posterior expected net host utility of each tier. Tier *j* is classified as viable if it belongs to the maximizing set for at least one signal; tied maxima are included with an absolute numerical tolerance of 10^*−*10^. This signalwise diagnostic allows the maximizing action to vary freely across signals and is distinct from the globally optimal monotone-policy calculation used in panels (a) and (c).

Figure **??**c compared synthetic systems with *K ∈ {* 3, 5, 6*}* . Host-benefit weights were evenly spaced, intervention burden was

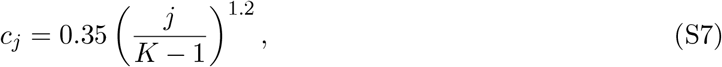

and the state prior is 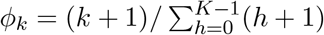. Panel (c) compares different state counts using this common prior-generation rule. Each system is optimized over its own monotone policy class. These parameterizations illustrate finite-policy geometry and piecewise-linear MIDL; they are not a controlled comparison of state resolution in one fixed population.

#### S2.3 Multi-segment aggregation illustration

Figure **??** illustrates Proposition 7.1 and Corollaries 7.1–7.2 using four intestinal segments and four functional guilds. The segment weights represent relative contributions to the modeled aggregate; they should not be interpreted as anatomical lengths or measured biomass fractions.

**Table S2.** Synthetic parameters used in the multi-segment aggregation illustration. Each composition *z*_*s*_ lies on the four-component probability simplex.

| Segment $s$ | $w_s$ | $\beta_s$ | $z_{s1}$ | $z_{s2}$ | $z_{s3}$ | $z_{s4}$ |
| --- | --- | --- | --- | --- | --- | --- |
| 1 | 0.10 | 0.75 | 0.55 | 0.25 | 0.15 | 0.05 |
| 2 | 0.15 | 0.55 | 0.35 | 0.35 | 0.20 | 0.10 |
| 3 | 0.25 | 0.35 | 0.18 | 0.32 | 0.30 | 0.20 |
| 4 | 0.50 | 0.15 | 0.08 | 0.22 | 0.35 | 0.35 |

The common reference profile and homogeneous baseline composition were

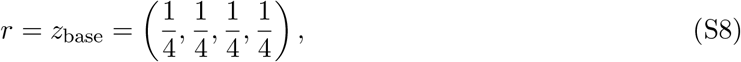

and the effective information-loss level was

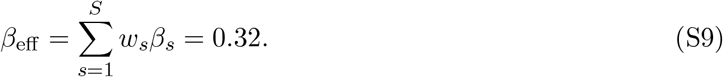

Compositional heterogeneity was varied through

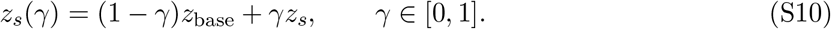

Thus, all segments have the same composition at *γ* = 0 and attain the profiles in Table S2 at *γ* = 1.

The interpolation in Eq. (S10) implies

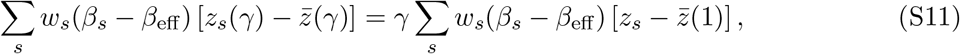

and therefore

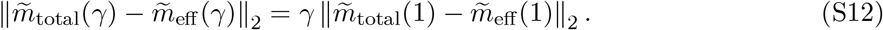

This linear dependence is specific to the chosen affine interpolation and should not be interpreted as a general law of intestinal heterogeneity.

#### S2.4 Ecological-screening illustration

Figure **??** combines a two-guild dynamical example with an expected-utility screening constraint. The parameters were selected to illustrate the transition from positive coexistence to a boundary equilibrium and the resulting restriction of the decision-feasible set. They were not estimated from empirical data.

**Table S3.** Parameters used in the ecological-screening illustration. All values are dimensionless and were selected for theoretical visualization.

| Panel | Quantity | Value |
| --- | --- | --- |
| (a) | Baseline growth vector | $\rho = (0.55, 0.80)^\top$ |
| | Intervention-response vector | $b = (-0.50, 0.55)^\top$ |
| | Interaction matrix | $A = \begin{pmatrix} 0.65 & 0.30 \\ 0.20 & 0.70 \end{pmatrix}$ |
| | Intervention range | $u \in [0, 1]$ |
| (b)–(c) | High-state prior probability | $\alpha = 0.60$ |
| | Reference signal distribution | $r = (1/2, 1/2)$ |
| | Aggressive-tier gross benefit | $b_1 = 1$ |
| | Aggressive-tier burden | $c_1 = \eta = 0.30$ |
| | Ecological reserve and burden | $\ell = \delta = (0, 1)$ |
| | Panel-(b) information loss | $\beta = 0.30$ |
| | Panel-(c) information-loss range | $\beta \in [0, 0.55]$ |
| | Ecological-margin floor | 38% of the identity-channel $U_E$ range |

##### S2.4.1 Two-guild dynamical model

Figure **??**a evaluates the gLV model in Eq. (35) at the parameters in Table S3. Local stability was evaluated using Eq. (37) with stability defined by Eq. (38). The crossing is *u*_crit_ *≈* 0.281553. The positive interior equilibrium is locally asymptotically stable for *u < u*_crit_, whereas the boundary equilibrium is locally asymptotically stable for *u > u*_crit_. The guild labels identify dynamical components only; no host-benefit ordering is inferred from these abundance equations. At *u* = *u*_crit_, one eigenvalue is zero and the two branches exchange stability through a transcritical bifurcation.

##### S2.4.2 Ecological constraint on the decision-feasible set

Figure **??**b–c use the binary-state channel and the three deterministic monotone policies defined above. Their convex hull is

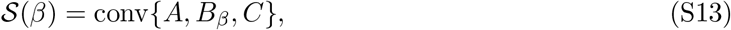

where the vertices are the closed-form policy outcomes given in the main text. Randomized mixtures of deterministic policies are represented by points within this convex hull.

The ecological-margin floor was defined relative to the identity-channel range:

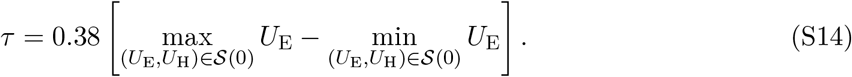

The factor 0.38 was selected to display both retained and excluded regions and is not intended as a biologically calibrated threshold. The admissible set is

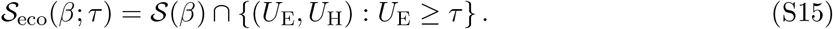

For each *β*, the unconstrained and constrained host-benefit frontiers were

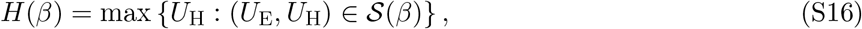

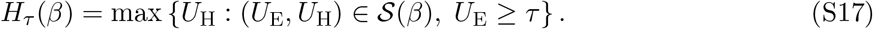

The ecological-screening gap was

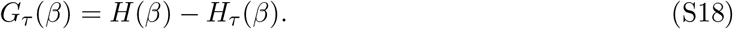

Because the optimization is performed over a convex hull, the constrained frontier may be attained by a randomized policy. Statewise, robust, or chance constraints would generally yield different admissible sets.

## References

Gregory P. Donaldson, S. Melanie Lee, and Sarkis K. Mazmanian. Gut biogeography of the bacterial microbiota. Nature Reviews Microbiology, 14(1):20–32, 2016. doi: 10.1038/nrmicro3552.

Jacob Folz, Rebecca Neal Culver, Juan Montes Morales, Jessica Grembi, George Triadafilopoulos, David A. Relman, Kerwyn Casey Huang, Dari Shalon, and Oliver Fiehn. Human metabolome variation along the upper intestinal tract. Nature Metabolism, 5(5):777–788, 2023. doi: 10.1038/s42255-023-00777-z.

Dari Shalon, Rebecca Neal Culver, Jessica A. Grembi, Jacob Folz, Peter V. Treit, Handuo Shi, Florian A. Rosenberger, Les Dethlefsen, Xiandong Meng, Eitan Yaffe, Andrés Aranda-Díaz, Philipp E. Geyer, Johannes B. Mueller-Reif, Sean Spencer, Andrew D. Patterson, George Triadafilopoulos, Susan P. Holmes, Matthias Mann, Oliver Fiehn, David A. Relman, and Kerwyn Casey Huang. Profiling the human intestinal environment under physiological conditions. Nature, 617(7961): 581–591, 2023. doi: 10.1038/s41586-023-05989-7.

David Blackwell. Equivalent comparisons of experiments. The Annals of Mathematical Statistics, 24 (2):265–272, 1953. doi: 10.1214/aoms/1177729032.

Dirk Bergemann, Benjamin Brooks, and Stephen Morris. The limits of price discrimination. American Economic Review, 105(3):921–957, 2015. doi: 10.1257/aer.20130848.

Alireza Fallah, Michael I. Jordan, Ali Makhdoumi, and Azarakhsh Malekian. The limits of price discrimination under privacy constraints, 2024.

Steven D. Allison and Jennifer B. H. Martiny. Resistance, resilience, and redundancy in microbial communities. Proceedings of the National Academy of Sciences, 105(Supplement 1):11512–11519, 2008. doi: 10.1073/pnas.0801925105.

Ashley Shade, Hannes Peter, Steven D. Allison, Didier L. Baho, Mercè Berga Helmut Bürgmann, David H. Huber, Silke Langenheder, Jay T. Lennon, Jennifer B. H. Martiny, Kristin L. Matulich, Thomas M. Schmidt, and Jo Handelsman. Fundamentals of microbial community resistance and resilience. Frontiers in Microbiology, 3:417, 2012. doi: 10.3389/fmicb.2012.00417.

David A. Relman. The human microbiome: ecosystem resilience and health. Nutrition Reviews, 70 (Supplement 1):S2–S9, 2012. doi: 10.1111/j.1753-4887.2012.00489.x.

Ronald A. Howard. Information value theory. IEEE Transactions on Systems Science and Cybernetics, 2(1):22–26, 1966. doi: 10.1109/TSSC.1966.300074.

Aharon Ben-Tal, Laurent El Ghaoui, and Arkadi Nemirovski. Robust Optimization. Princeton University Press, Princeton, 2009. doi: 10.1515/9781400831050.

Arkadi Nemirovski and Alexander Shapiro. Convex approximations of chance constrained programs. SIAM Journal on Optimization, 17(4):969–996, 2006. doi: 10.1137/050622328.

Richard R. Stein, Vanni Bucci, Nora C. Toussaint, Charlie G. Buffie, Gunnar Rätsch, Eric G. Pamer, Chris Sander, and João B. Xavier. Ecological modeling from time-series inference: insight into dynamics and stability of intestinal microbiota. PLoS Computational Biology, 9(12):e1003388, 2013. doi: 10.1371/journal.pcbi.1003388.

Ophelia S. Venturelli, Alex V. Carr, Garth Fisher, Ryan H. Hsu, Rebecca Lau, Benjamin P. Bowen, Susan Hromada, Trent Northen, and Adam P. Arkin. Deciphering microbial interactions in synthetic human gut microbiome communities. Molecular Systems Biology, 14(6):e8157, 2018. doi: 10.15252/msb.20178157.

Katharine Z. Coyte, Jonas Schluter, and Kevin R. Foster. The ecology of the microbiome: networks, competition, and stability. Science, 350(6261):663–666, 2015. doi: 10.1126/science.aac4259.

John Aitchison. The Statistical Analysis of Compositional Data. Chapman and Hall, London, 1986. doi: 10.1007/978-94-009-4109-0.

Gregory B. Gloor, Jean M. Macklaim, Vera Pawlowsky-Glahn, and Juan J. Egozcue. Microbiome datasets are compositional: and this is not optional. Frontiers in Microbiology, 8:2224, 2017. doi: 10.3389/fmicb.2017.02224.

